# Modular organization of the murine locomotor pattern in presence and absence of sensory feedback from muscle spindles

**DOI:** 10.1101/470492

**Authors:** Alessandro Santuz, Turgay Akay, William P. Mayer, Tyler L. Wells, Arno Schroll, Adamantios Arampatzis

## Abstract

For exploiting terrestrial and aquatic locomotion, vertebrates must build their locomotor patterns based on an enormous amount of variables. The great number of muscles and joints, together with the constant need for sensory feedback information (e.g. proprioception), make the task of creating and controlling movement a problem with overabundant degrees of freedom. It is widely accepted that the central nervous system might simplify the creation and control of movement. This could happen through the generation of activation patterns, which are common to many different muscles, rather than specific to individual muscles. These activation patterns, called muscle synergies, can be extracted from electromyographic data and describe the modular organization of movement. We extracted muscle synergies from the hindlimb muscle activities of wild type and genetically modified mice, in which sensory feedback from muscle spindles is eliminated. Muscle spindle-deficient mice underwent a modification of the temporal structure (motor primitives) of muscle synergies that resulted in diminished functionality during walking. In addition, both the temporal and spatial components (motor modules) of muscle synergies were severely affected when external perturbations were introduced of when animals were immersed in water. These findings show that group Ia/II sensory feedback from muscle spindles regulates motor function in normal and perturbed walking. Moreover, when group Ib Golgi tendon organ feedback is lacking due to the reduction of gravitational load in conditions of enhanced buoyancy, the modular organization of swimming is almost completely compromised.

**Significance statement:** Locomotion on land and in water requires the coordination of a great number of muscle activations and joint movements. Moreover, constant feedback about the position of own body parts in relation to the surrounding environment and the body itself (proprioception) is required to maintain stability and avoid failure. The theory of muscle synergies states that the central nervous system might control muscles in orchestrated groups (synergies) rather than individually. We used this concept on genetically modified mice, lacking one of the two classes of proprioceptors. Our results provide evidence that proprioceptive feedback is required by the central nervous system to accurately tune the modular organization of locomotion.

## Introduction

In order to move through different environments, vertebrates must coordinate an overwhelming number of muscles and joints. The complex task of generating and controlling movement is achieved by a fine interplay of muscle activations, originated from and tuned by the brain and spinal cord (i.e. the central nervous system, CNS). Sensory feedback to the CNS plays a significant role in influencing the control of movement patterns. For instance, the ability to sense the position of own body parts in relation to the surrounding environment and the body itself, called proprioception, is of clear importance (1). During locomotion, the CNS continuously elaborates the information coming from proprioceptors, a myriad of sensors (2) located in muscles (group Ia/II muscle spindles) and tendons (group Ib Golgi tendon organs, GTOs). It is well known that the loss of proprioceptive feedback may severely affect the motor function in mammals and is a symptom commonly associated with numerous neurological and orthopaedic conditions (3–8). There is also evidence that aging results in the decline of proprioceptors, especially those located in the more distal portions of the limbs (9). Yet, the mechanisms underlying the capability of the CNS to integrate proprioceptive signals for tuning motor coordination are still obscure (6, 10).

Since the first structured attempts of describing motor coordination (11), how animals can achieve accurate, rapid movements choosing from an enormous amount of muscle/joint configurations has been an open question. A common theoretical framework for describing the creation and control of movement has been successfully used in humans and other animals in the past three decades: the concept of muscle synergies (12–16). It is suggested that the CNS might not be controlling each muscle and joint separately. Instead, movement control might be simplified by the synchronous activation of selected groups of muscles, in patterns called synergies (17). Time-dependant electromyographic (EMG) activities recorded from many muscles at the same time can be conveniently represented in a combination of time-independent weights, also called motor modules (18, 19), and time-dependent coefficients, also called motor primitives (14, 19). Motor modules and motor primitives are, together, a compact representation of the modular organization of the locomotor pattern.

In this study we assessed the role of proprioceptive sensory feedback in the modular organization of the murine locomotor pattern. We examined a mouse line in which a mutation in the *Egr3* (early growth response 3) gene selectively impairs group Ia/II muscle spindles, eliminating the feedback from one of the two classes of proprioceptors (20). We extracted muscle synergies from the EMG activity recorded from the lower limb of mice during three tasks: treadmill walking, swimming and “cautious” treadmill walking. The first two tasks differed in the contribution of input from group Ib sensory afferents supplying GTOs (6). With the third task we applied electrical stimulation of the saphenous nerve at regular intervals in order to elicit a cautious way of walking between two consecutive stimulations (8). We extracted synergies using non-negative matrix factorization (NMF) (21, 22). We quantified the dynamic stability (23) of walking by calculating the maximum finite-time Lyapunov exponent (MLE) of the hip, knee and ankle joint angles (24, 25).

Our results show that the modular organization of locomotion in mice is continuously tuned by proprioceptive feedback. The absence of feedback from muscle spindles impairs the dynamic stability and the temporal component (i.e. motor primitives) of muscle synergies during normal walking and both the temporal and spatial components (i.e. motor modules) during cautious walking. This results in a) moderate impairment of normal walking and b) the inability of the motor system to cope with external expected perturbations. The absence of feedback from muscle spindles and GTOs, made possible by the immersion of animals in water, creates a strong impairment of both temporal and spatial elements of muscle synergies, resulting in an almost complete loss of function. These findings show that, in mice, muscle spindles and GTOs afferents provide important information to the CNS for correctly tuning the modular organization of locomotion.

## Results

### Locomotion parameters

We first compared the basic parameters of locomotion in WT and *Egr3^−/−^* mice of GR1. The stance duration in walking did not differ significantly (p = 0.137) between WT (192 ± 30 ms) and *Egr3^−/−^* (178 ± 22 ms). The swing times in walking did not differ significantly as well (p = 0.140) between WT (100 ± 21 ms) and *Egr3^−/−^* (91 ± 13 ms). Cadence was also not influenced by the absence of muscle spindles (419 ± 59 steps/min for WT and 447 ± 34 steps/min for *Egr3^−/−^* mice, p = 0.100). In swimming, the locomotion cycle duration was significantly larger (p = 0.007) in *Egr3^−/−^* (243 ± 57 ms) compared to WT (181 ± 33 ms). Consequently, the cadence decreased, from 682 ± 109 (WT) to 528 ± 165 (*Egr3^−/−^*) cycles/min (p = 0.020). Then we looked into the basic parameters of locomotion in WT and *Egr3^−/−^* mice of GR2, comparing walking on treadmill during normal walking and cautious (with electrical stimulation of the saphenous nerve walking. The stance duration in WT did not differ significantly (p = 0.312) between normal (144 ± 8 ms) and cautious walking (136 ± 11 ms). The swing times did differ significantly (p = 0.029, 101 ± 8 ms during normal and 152 ± 18 ms during cautious walking). Cadence decreased when switching from normal (493 ± 24 steps/min) to cautious walking (420 ± 39 steps/min, p = 0.025). In *Egr3^−/−^* mice, the stance duration was almost identical (p = 0.997) in normal (130 ± 7 ms) or cautious walking (130 ± 17 ms), while the swing differed significantly (p = 0.001, 80 ± 11 ms in normal and 114 ± 11 ms in cautious walking). Cadence significantly decreased (from 573 ± 27 to 493 ± 38 steps/min, p = 0.006) in cautious walking. In absence of proprioceptive feedback form the muscle spindles, the basic timing parameters of walking were not affected. However, the average duration of the swimming cycle increased. Moreover, both in WT and *Egr3^−/−^* mice, occasional perturbation of walking through the electrical stimulation of the saphenous nerve, causing a more cautious walking (see methods), produced an increase in the swing duration and a decrease in cadence, while stance was unaffected.

### EMG

We have next compared the activity patterns of flexor and extensor muscles of GR1 mice to investigate the effect of muscle spindle removal on locomotor pattern generation. Figure 1 is a representation of the average recorded EMG activities for both WT and *Egr3^−/−^* during walking and swimming. One locomotor cycle is represented as circular plots, whereas the EMG activities, normalized to the maximum activity of each muscle, are coded by the darkness of the color. That is, darker color indicates more activity. In the absence of the muscle spindles, EMG activity pattern during walking is subtly changed, most prominently as a prolonged TA activity at the end of swing phase. Furthermore, during swimming, flexor muscles are sequentially activated beginning with the most proximal hip flexor muscle (IP) and propagating distally towards the ankle flexor TA in the wild type. In contrast, in *Egr3^−/−^* mice the flexor muscles are activated simultaneously. This data suggests that, in absence of proprioceptive sensory feedback, locomotor pattern is still functional with subtle differences, but is severely affected when proprioceptive from both the muscle spindles and the Golgi tendon organs are missing such as during *Egr3^−/−^* swimming (6).

**Fig. 1.**
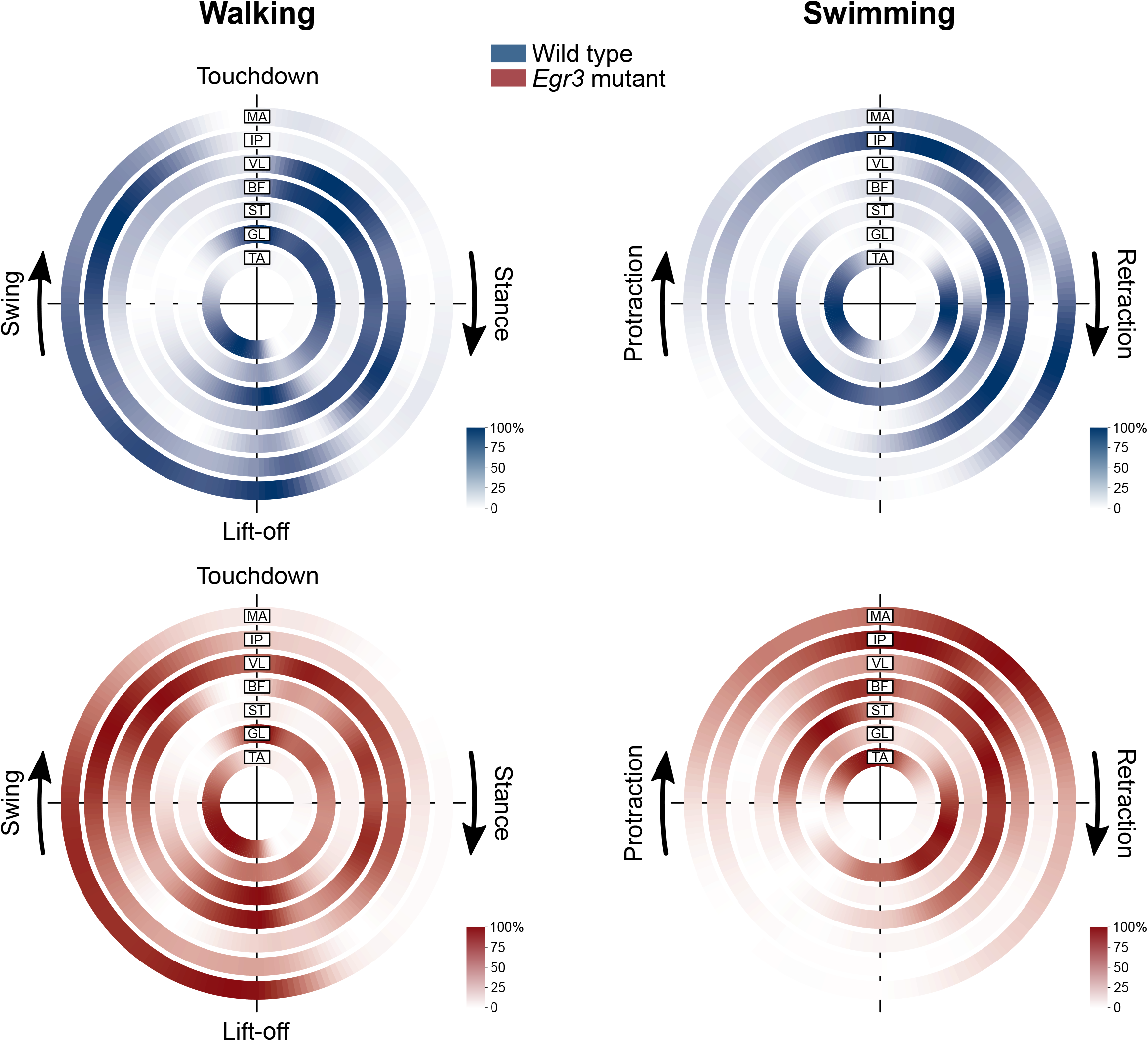
Average EMG patterns represented as donut plots for walking and swimming in wild type and mutant animals (GR1). The duration of stance and swing in walking are represented with the same amount of time points for better visualization. Color scales go from minimum activation (white) to muscle-specific maximum recorded activation (blue for wild type and red for *Egr3* mutant). Muscle abbreviations: MA=*gluteus maximus*, IP=*iliopsoas*, VL=*vastus lateralis*, BF=*biceps femoris*, ST=*semitendinosus*, GL=*gastrocnemius lateralis*, TA=*tibialis anterior*.

### Local dynamic stability

We investigated the effect of muscle spindle removal on the local dynamic stability during treadmill walking (animals of GR1). The mean MLE values are reported in Table 1. These values showed an increase of the MLE from the most proximal (hip) to the most distal (ankle) joint (see Figure 2). Analyzing the differences between the MLE values of the three joints and the respective confidence intervals, we could confirmed the significance of this increase (0.0949 [0.0608, 0.1273], 0.1196 [0.0842, 0.1584] when taking the differences between the MLE of the knee and the hip of WT and *Egr3^−/−^* mice, respectively; 0.1628 [0.1364, 0.1883], 0.1276 [0.0955, 0.1586] when taking the differences between the MLE of the ankle and the knee of WT and *Egr3^−/−^* mice, see Figure 2). Furthermore, we could detect significantly higher MLE values for hip, knee, and ankle joints in the *Egr3^−/−^* mice than in WT mice (Table 1 and Figure 2). Confidence intervals for the MLE difference of population means in WT and *Egr3^−/−^* were [0.0008; 0.0063], [0.0011; 0.0066] and [0.0007; 0.0060] for the hip, knee and ankle joint, respectively. The higher values for *Egr3^−/−^* compared to WT indicated a lower local dynamic stability for the mutant mice in each limb joint. This difference has to be considered as significant since zero is lying outside the confidence intervals. Confidence intervals for the unbiased effect size estimator Hedges’ g were [0.2058; 1.6172], [0.2487; 1.6405] and [0.1736; 1.5421] with means 0.88, 0.94 and 0.86 for the hip, knee and ankle joint, indicating a strong effect for all three joints. This data shows that the dynamic stability of the hip, knee and ankle joint is affected by the removal of muscle spindles, condition that produces an increased instability of the system.

**Table 1.** Mean value, difference and confidence intervals in local dynamic stability as well as unbiased effect size estimates (Hedges’ g) between *Egr3* mutant and wild type mice for hip, knee and ankle angle. Positive differences (Δ_W,M_ > 0) denote bigger values in the mutants and though greater instability, whereas negative differences imply the contrary. The Hedges’ g effect size shows the bias corrected standardized differences between mutant and wild type means. Asterisks highlight the 95% confidence intervals (CI) which do not contain zero and therefore have to be interpreted as significant.

| Local dynamic stability |  |  |  |  |  |  |  |  |
| --- | --- | --- | --- | --- | --- | --- | --- | --- |
| Joint | MLE WT | 95% CI | MLE KO | 95% CI | $\Delta_{W,M}$ | 95% CI | $g$ | 95% CI |
| Hip | 1.1435 | [1.0256, 1.2782]* | 1.3968 | [1.2563, 1.5484]* | 0.2532 | [0.0008, 0.0063]* | 0.88 | [0.2058, 1.6172]* |
| Knee | 1.2385 | [1.1183, 1.3697]* | 1.5163 | [1.3730, 1.6732]* | 0.2779 | [0.0746, 0.4622]* | 0.94 | [0.2487, 1.6405]* |
| Ankle | 1.4013 | [1.2877, 1.5264]* | 1.6439 | [1.5124, 1.7906]* | 0.2395 | [0.0514, 0.4190]* | 0.86 | [0.1736, 1.5421]* |

**Fig. 2.**
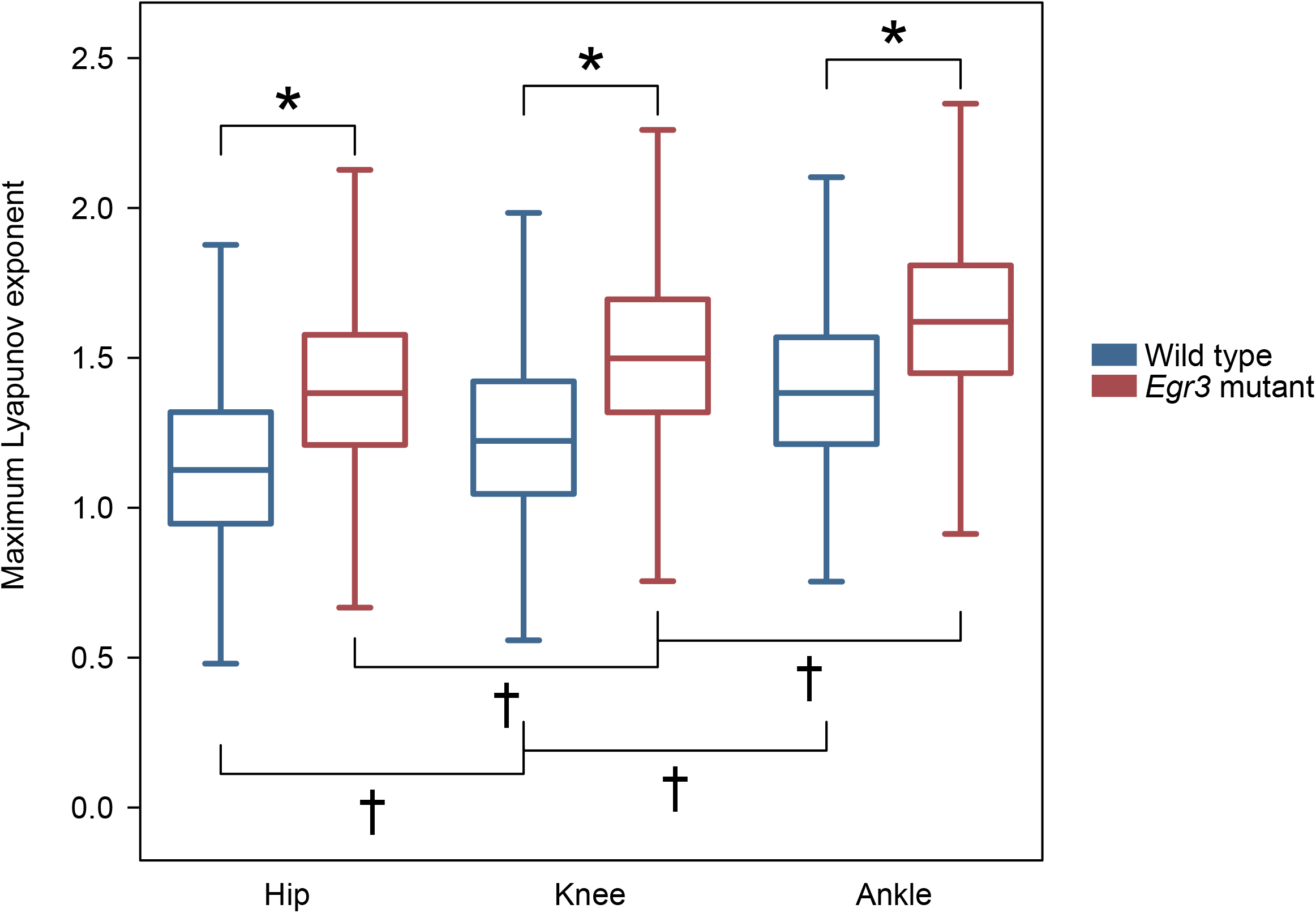
Box plots representing the maximum Lyapunov exponent (MLE) values of the three joint angles for wild type and *Egr3* mutant bootstrapped individuals during walking. A faster divergence, represented by bigger MLE values, indicates worse dynamic stability. Asterisks represent significant differences between groups, obtained from the analysis of confidence intervals. Similarly, daggers denote significant differences between joints.

### Muscle synergies

We extracted muscle synergies from the EMG data recorded during walking (GR1 and GR2) and swimming (GR1). The minimum number of synergies which best accounted for the EMG data variance (i.e. the factorization rank) of GR1 mice was, in walking, 3.3 ± 0.5 (WT) and 3.0 ± 0.3 (*Egr3^−/−^*); in swimming, the values were of 3.7 ± 0.5 (WT) and 2.6 ± 0.6 (*Egr3^−/−^*). In GR2, the values for walking were of 3.1 ± 0.3 (WT, normal walking), 3.1 ± 0.4 (WT, cautious walking), 3.0 ± 0.5 (*Egr3^−/−^*, normal walking) and 3.3 ± 0.6 (*Egr3^−/−^*, cautious walking). Values represent the mean and standard deviation of the 10000 runs per condition (Figure 3 and Figure 4).

In walking, the first synergy functionally referred to the body weight acceptance and propulsion, with a major involvement of the knee extensors and flexors and of the ankle extensors. The second synergy described the early swing phase, to which the hip rotators, knee flexors and ankle flexors mainly contributed. The third synergy identified the late swing, showing the involvement of the hip rotators and flexors and of the ankle flexors. In swimming, the first and second synergies functionally referred to the retraction phase. In WT, the hip flexors mainly contributed to the first synergy, while all the other muscles with the exception of the ankle flexors were involved in the second synergy. In *Egr3^−/−^* animals, the first synergy showed the additional contribution of the ankle flexors, while the second synergy was highlighting the involvement of only the hip rotators and the knee flexors and extensors. The third and last synergy was associated with the late retraction phase of swimming, where knee flexors and ankle flexors mainly contributed in WT animals. In *Egr3^−/−^* animals, this last synergy was mainly associated with the activity of knee flexors and extensors and of the ankle extensors. Three fundamental synergies were extracted from the EMG activity of six or seven muscles regardless of the condition (normal walking, cautions walking, or swimming in WT or *Egr3^−/−^* mice), each synergy functionally corresponded to a specific task necessary for locomotion.

**Fig. 3.**
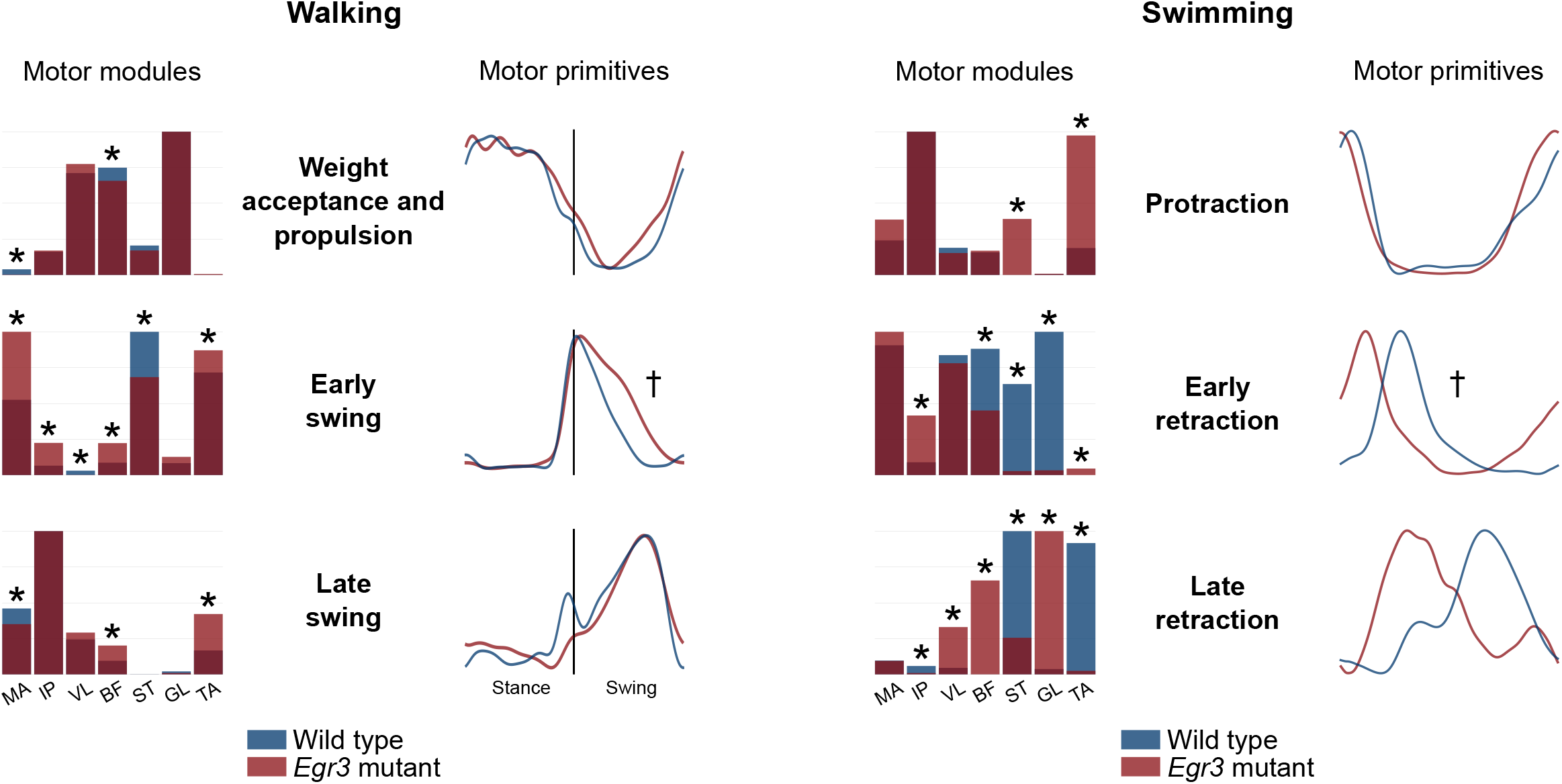
Average motor modules and motor primitives of the three fundamental synergies for walking and swimming in wild type and *Egr3* mutant animals (group GR1). In walking the first synergy referred to the weight acceptance and propulsion, the second synergy to the early swing and the third synergy to the late swing phase. In swimming, the first synergy described the protraction, the second the early retraction and the third the late retraction phase. The motor modules are presented on a normalized y-axis base, with asterisks denoting significant differences obtained by the estimated 95% confidence intervals. For the motor primitives, the x-axis full scale represents one locomotion cycle (in walking, stance and swing were normalized to the same amount of points and divided by a vertical line) and the y-axis the normalized amplitude, with daggers denoting an increased FWHM. Muscle abbreviations: MA=*gluteus maximus*, IP=*iliopsoas*, VL=*vastus lateralis*, BF=*biceps femoris*, ST=*semitendinosus*, GL=*gastrocnemius lateralis*, TA=*tibialis anterior*.

In order to deepen our analysis of the modular organization of locomotion in presence and absence of muscle spindles, we investigated motor modules and motor primitives separately. The assessment of the temporal parameters of the motor primitives in WT and *Egr3^−/−^* mice of GR1 revealed specific changes in the FWHM and CoA in absence of proprioceptive feedback form muscle spindles. The FWHM of the motor primitives was increased (widening) in mutants in the second synergy of both walking (early swing) and swimming (early retraction, see Table 2). The CoA of motor primitives shifted later in time in all synergies of walking and earlier in the early retraction and late retraction synergy of swimming (Table 2). Other differences could not be included in the 95% confidence interval estimated from the bootstrapped data. This data suggests that the sequence of motor primitives during walking is subtly changed in the absence of muscle spindle feedback but severely compromised during swimming when feedback from the GTOs is strongly reduced.

Investigation of motor modules of GR1 mice revealed more severe changes than the motor primitives. Motor modules in walking showed a significant difference in the contribution of MA and BF muscles in the first (weight acceptance and propulsion) synergy. In the early swing synergy, the muscles MA, IP, VL, BF, ST and TA all proved to be used differently in *Egr3^−/−^* mice compared to WT. The late swing synergy only showed a difference contribution of MA, BF and TA muscles (see asterisks in Figure 3). In swimming, the motor modules of the protraction synergy only differed in the contribution of ST and TA muscles, while the synergy for early retraction included more differences, with only MA and VL retaining the same weight. The modules of the late retraction synergy showed a significant different contribution of all muscles but the MA (see asterisks in Figure 3). These outcomes highlight that, together with the changes observed in the motor primitives, the time-invariant modular organization of motion was affected by the absence of muscle spindles. However, changes were more dramatic in swimming than in walking.

**Table 2.** The motor primitives’ full width at half maximum (FWHM) and center of activity (CoA) are reported for group GR1 as percentage differences between wild type (W) and *Egr3* mutants (M) values (Δ_W,M_ ± standard deviation). Positive differences (Δ_W,M_ > 0) denote bigger values in the mutants, whereas negative differences imply the contrary. Asterisks highlight the 95% confidence intervals (CI, based on actual FWHM and CoA values) which do not contain the zero. The coefficient of determination (R^2^) indicates the average similarity of motor primitives between wild type and mutants.

|  |  | Motor primitives |  |  |  |  |
| --- | --- | --- | --- | --- | --- | --- |
| | | $R^2$ | FWHM | | CoA | |
| | | Values | $\Delta_{W,M}$ | 95% CI | $\Delta_{W,M}$ | 95% CI |
| Walking | Weight acceptance and propulsion | 0.79<br>$\pm 0.02$ | -3.0%<br>$\pm 2.4\%$ | [-6.4, +1.4] | +22.5%<br>$\pm 3.8\%$ | [+9.0, +18.0]* |
| | Early swing | 0.42<br>$\pm 0.14$ | +18.8%<br>$\pm 3.3\%$ | [+8.2, +16.7]* | +9.2%<br>$\pm 1.9\%$ | [+7.2, +16.5]* |
| | Late swing | 0.55<br>$\pm 0.05$ | -9.0%<br>$\pm 2.8\%$ | [-10.6, -2.6]* | +14.0%<br>$\pm 1.8\%$ | [+16.9, +27.8]* |
| Swimming | Protraction | 0.72<br>$\pm 0.04$ | -8.0%<br>$\pm 3.3\%$ | [-9.4, -0.9]* | +14.6%<br>$\pm 9.8\%$ | [-7.6, +61.7] |
| | Early retraction | -0.74<br>$\pm 0.11$ | +19.5%<br>$\pm 3.1\%$ | [+8.6, +16.5]* | -35.9%<br>$\pm 5.9\%$ | [-29.7, -15.2]* |
| | Late retraction | -1.16<br>$\pm 0.10$ | +6.5%<br>$\pm 3.8\%$ | [-0.6, +10.7] | -39.6%<br>$\pm 2.4\%$ | [-61.0, -47.8]* |

Following the same approach, we then analyzed motor primitives and motor modules of both WT and *Egr3^−/−^* animals during normal and cautious walking (GR2). In WT mice, cautious walking was related to a widening in the first and third primitives (weight acceptance/propulsion and late swing, Table 3 and Figure 4). Moreover, also the CoA underwent a shift earlier in time in the same synergies (Table 3 and Figure 4). However, this was not the case in *Egr3^−/−^* animals, where the FWHM did not increase in any primitive and the CoA did not show any changes (Table 3 and Figure 4). Additionally, motor modules differed in both WT and mutants in the first two synergies, with only a very limited amount of muscles responsible for the changes. This additional data shows that introducing an external perturbation during treadmill walking does not affect the general modular organization of locomotion. However, the perturbation has strong effects on the timing of the locomotor output, specifically widening two out of three motor primitives. Interestingly, this behavior could only be detected in WT animals, suggesting that muscle spindle-deficient mice were not able to mitigate the effects of the perturbation by introducing adjustments in the locomotor patterns.

**Table 3.** The motor primitives’ full width at half maximum (FWHM) and center of activity (CoA) are reported for group GR2 as percentage differences between normal and cautious (with electrical stimulation of the saphenous nerve) walking in both wild type (W) and *Egr3* mutants (M) values (ΔW,M ± standard deviation). Positive differences (Δ_W,M_ > 0) denote bigger values in cautious walking, whereas negative differences imply the contrary. Asterisks highlight the 95% confidence intervals (CI) which do not contain the zero. The coefficient of determination (R^2^) indicates the average similarity of motor primitives between normal and cautious walking.

|  |  | Motor primitives |  |  |  |  |
| --- | --- | --- | --- | --- | --- | --- |
| | | $R^2$ | FWHM | | CoA | |
| | | Values | $\Delta W_{W,M}$ | 95% CI | $\Delta W_{W,M}$ | 95% CI |
| Wild type | Weight acceptance and propulsion | 0.53<br>$\pm 0.14$ | +11.4%<br>$\pm 5.7\%$ | [+0.3%, +17.6%]* | -17.1%<br>$\pm 7.4\%$ | [-20.9%, -1.5%]* |
| | Early swing | 0.92<br>$\pm 0.03$ | -9.6%<br>$\pm 5.9\%$ | [-12.5%, +1.1%] | -1.7%<br>$\pm 2.2\%$ | [-7.9%, +3.1%] |
| | Late swing | 0.72<br>$\pm 0.06$ | +22.4%<br>$\pm 4.8\%$ | [+7.4%, +18.0%]* | -20.0%<br>$\pm 5.2\%$ | [-57.1%, -19.7%]* |
| *Egr3*<sup>-/-</sup> | Weight acceptance and propulsion | 0.82<br>$\pm 0.05$ | -15.2%<br>$\pm 4.1\%$ | [-20.9%, -6.4%]* | +4.0%<br>$\pm 7.1\%$ | [-5.1%, +9.4%] |
| | Early swing | 0.86<br>$\pm 0.05$ | -8.5%<br>$\pm 6.3\%$ | [-14.0%, +2.4%] | +0.3%<br>$\pm 3.0\%$ | [-7.2%, +7.9%] |
| | Late swing | 0.57<br>$\pm 0.09$ | +1.6%<br>$\pm 5.9\%$ | [-8.6%, +9.7%] | +2.2%<br>$\pm 3.5\%$ | [-7.9%, +15.2%] |

**Fig. 4.**
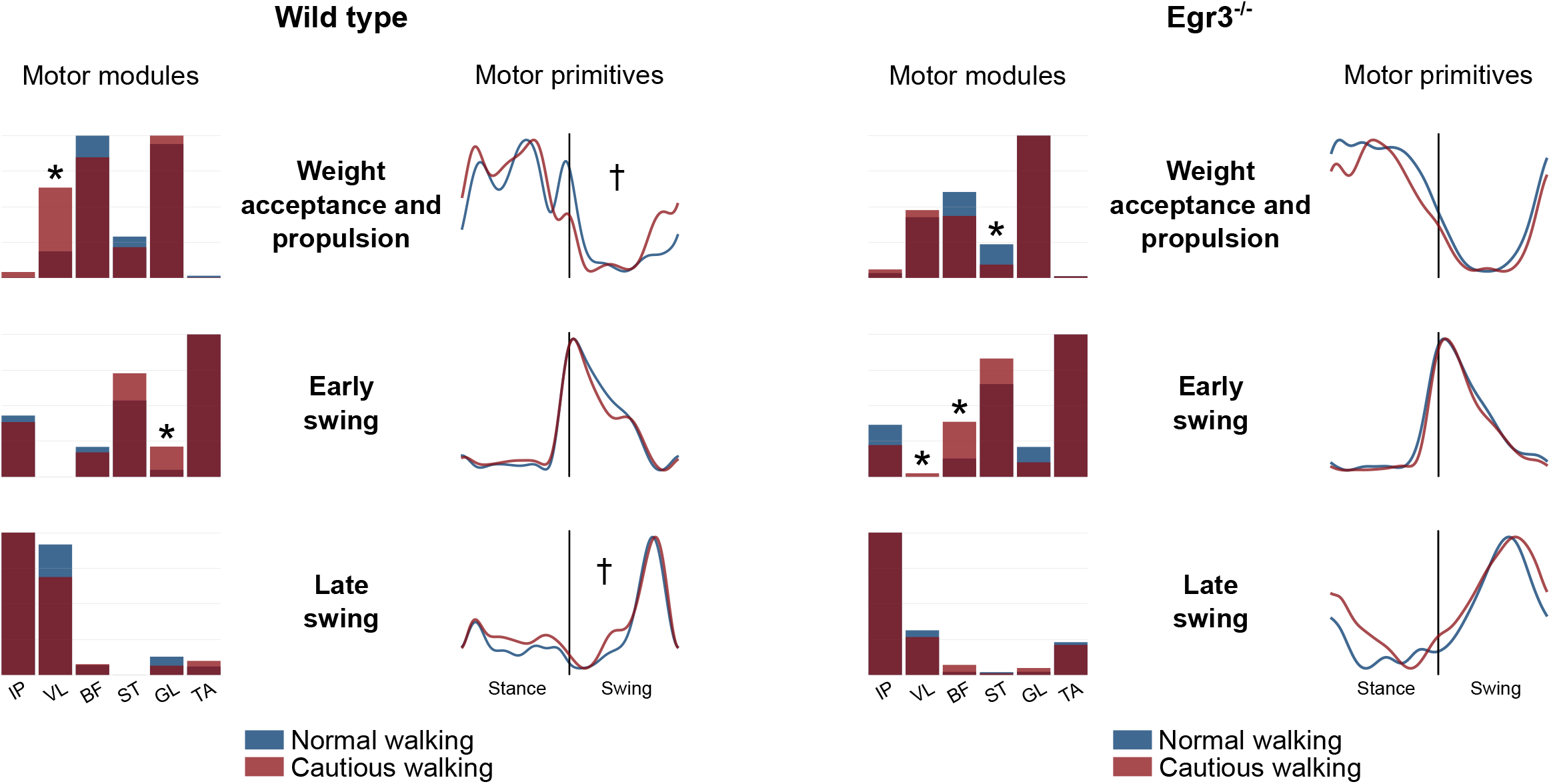
Average motor modules and motor primitives of the three fundamental synergies for normal and cautious walking (see methods) in wild type and *Egr3* mutant animals (group GR2). The first synergy referred to the weight acceptance and propulsion, the second synergy to the early swing and the third synergy to the late swing phase. The motor modules are presented on a normalized y-axis base, with asterisks denoting significant differences obtained by the estimated 95% confidence intervals. For the motor primitives, the x-axis full scale represents one locomotion cycle (stance and swing were normalized to the same amount of points and divided by a vertical line) and the y-axis the normalized amplitude, with daggers denoting an increased FWHM. Muscle abbreviations: IP=*iliopsoas*, VL=*vastus lateralis*, BF=*biceps femoris*, ST=*semitendinosus*, GL=*gastrocnemius lateralis*, TA=*tibialis anterior*.

## Discussion

In this study, we analyzed the neuromuscular control of locomotion in adult *Egr3^−/−^* (in which all muscle spindles are ablated and GTOs are left intact) and WT mice. We hypothesized that proprioceptive feedback from muscle spindles is necessary for generating an accurately tuned locomotor pattern and maintaining dynamic stability during locomotion. We predicted a lower dynamic stability in *Egr3^−/−^* compared to WT mice, together with a loss in temporal accuracy of the neuromuscular control of treadmill walking and swimming. The higher instability in *Egr3^−/−^* mice was confirmed by higher MLE of the hindlimb joint angles. We could also confirm, in muscle spindle-deficient animals, a decreased temporal accuracy of motor control (i.e. a widening in the temporal structure, described by the motor primitives) in one out of three muscle synergies in both walking and swimming and clear time-shifts in almost all synergies during walking and swimming. The widening of motor primitives was also visible when comparing normal and cautious (see methods) walking in WT mice, but not in *Egr3^−/−^*, suggesting that muscle spindles are crucial to accurately tune the timing of locomotion patterns. Furthermore, it was evident that both walking and swimming shared the same number of muscle synergies. This feature did not differ between *Egr3^−/−^* and WT animals, indicating the use of a consistent set of neural elements for controlling locomotion both with and without proprioceptive sensory feedback from muscle spindles.

### Muscle spindle feedback is required for stable locomotion

The MLE quantify how a dynamical system responds to small perturbations (26, 27), revealing its ability to maintain stability (24, 28). Increased MLE correspond to a more chaotic and unstable dynamical system (29, 30). In humans, previous studies found an increase in the MLE during walking in patients with focal cerebellar lesion (31) and with moderate neurological gait disorders (32). We also showed that increased MLE is detectable when inexperienced barefoot runners transition from shod to barefoot running (33) and when healthy adults walk and run over uneven surfaces (34). Similarly, during murine gait, it can be expected that the absence of feedback from muscle spindles introduce input errors (35). In other words, when muscle spindles are missing, animals might not be able to retrieve accurate information about the state of the system. In fact, we found that *Egr3^−/−^* mice showed higher MLE (i.e. lower stability) than the WT. This finding suggests that an increase in input errors (35) resulted in reduced stability in muscle spindle-deficient mice during treadmill walking.

Moreover, when comparing WT and *Egr3^−/−^* mice, we found the MLE to be lower in the proximal (hip) than in the distal (ankle) joints during walking. This characteristic (i.e. greater instability in distal than in proximal joints) has been previously found also in young and older humans (36). Mechanically attributed to the greater inertia of the trunk compared to the limbs (36), it has been interpreted as a reflection of the nervous system’s need to prioritize the stability of head and trunk over other inferior segments (37).

Put together, the present outcomes suggest that the absence of feedback from muscle spindles in mice is a source of internal perturbation that hinders the retrieval of accurate information about the state of the system. This results in a decreased stability of *Egr3^−/−^* compared to WT mice.

### Proprioceptive feedback affects spatial and temporal structure but not the number of synergies

Literally, synergy means “working together” (from the Greek συνεργóς). In the past few decades, many authors suggested that the control of movement might happen through the combination of a small set of neurophysiological entities named muscle synergies, rather than through the activation of muscles individually (12, 14, 38, 39). Muscle synergies for locomotion are remarkably retained across mammals, for instance in humans (14, 15) and rats (16), where two to five synergies are usually enough to describe locomotion, depending on the number of recorded muscle activities and on the stage of development. In our study, the minimum number of synergies necessary to reconstruct the original EMG signal in WT mice was three in both walking and swimming. This outcome did not change in *Egr3^−/−^* mice, indicating that the number of fundamental neural control elements did not change across locomotion types and was not affected by the absence of muscle spindles. However, when comparing the effects of muscle spindle ablation on locomotor patterns, swimming was more affected than walking. In fact, functionality was quite preserved during walking, that is mice could advance on the treadmill. Conversely, the functionality of the locomotor pattern during swimming was severely compromised. Only two out of 17 *Egr3^−/−^* mice could move in the water, although leg movements were clearly ataxic as previously described (6). The remaining mice moved their legs in a highly ataxic manner unable to provide propulsion. Regardless of whether the *Egr3^−/−^* mice locomote in water, the temporal structure of the locomotor pattern was severely compromised, although the number of fundamental neural control elements was preserved. It has been previously shown that mice lacking proprioceptive sensory neurons from muscle spindles and GTOs (Pv::cre;Isl2::DTA) can neither walk, nor swim (6). In *Egr3^−/−^* animals, GTOs are intact instead and contribute, during walking, to provide some sensory feedback. When animals are immersed in water, however, also the contribution from GTOs is strongly reduced due to the lower gravitational load induced by buoyancy. This often results in ataxic movements (6) and is likely the main reason for the aberrant locomotor pattern that we observed. These observations clearly indicate that proprioception plays an important role in the global sensory weighting during both swimming and walking. However, the number of fundamental neural control elements (i.e. muscle synergies) is not affected by the absence of proprioceptive feedback.

To further analyze the effect of muscle spindle deficiency on the modular organization of motion, we assessed the similarity between motor primitives of *Egr3^−/−^* and WT mice using the coefficient of determination R^2^. In previous studies we showed that, in humans, typical R^2^ values between trials of the same condition (i.e. repeatability without electrode removal) can range from 0.75 to 0.95 when comparing motor primitives (19, 34, 40). In our study, it was evident from the values presented in Table 2 that only the R^2^ of the first synergy in both walking (weight acceptance and propulsion) and swimming (protraction) was very close to the repeatability values found in humans (19, 34, 40). All the other synergies showed lower values when comparing mutant and WT animals, suggesting that the shape of the basic activation patterns was influenced by the selective removal of muscle spindles. Two metrics for evaluating the shape of motor primitives are the FWHM and CoA, indicators of duration and timing, respectively (34, 41, 42). We found an increased FWHM (which corresponds to a widening) in the early swing synergy of walking and in the early retraction synergy of swimming. Contextually, we found a reduction in the FWHM (narrowing) in the late swing primitive of walking and in the protraction primitive of swimming. In addition, the CoA increased in all the synergies of walking in mutants, denoting a shift of motor primitives later in time. Conversely, the two retraction primitives of swimming underwent, in mutants, a clear shift earlier in time, evidenced by the lower CoA. These observations confirm an alteration in the time structure and shape of motor primitives in the motor control of *Egr3^−/−^* mice. The absence of proprioceptive feedback from muscle spindles in *Egr3^−/−^* mice has previously been reported to be a source of internal perturbation for locomotor patterns (6). Changes in the shape and timing of motor primitives reflect the diminished functionality during walking and swimming when motor control is impaired by the absence of proprioceptive feedback.

Motor modules explain the relative contribution (or weight) of individual muscles within a specific synergy. We found that the motor modules of the first synergy in both walking (weight acceptance and propulsion) and swimming (protraction) were influenced by the lack of muscle spindles. Only a few muscles contributed to this adjustment due to the absence of muscle spindles: the hip rotators (MA muscle), hip extensors/knee flexors in walking (BF muscle) and the hip extensors/knee flexors (ST muscle) and ankle flexors (TA muscle) in swimming. Instead, the second synergy (early swing in walking and early retraction in swimming) showed an increased number of adjustments in the relative contributions of muscles to both locomotion types (see asterisks in Figure 3 for details). The third synergy for walking (late swing) highlights the higher contribution of BF and TA and a reduced contribution of MA. These adjustments in the third synergy during walking explain the exaggerated foot lifting of mutants compared to WT during walking (6). In swimming, the late retraction synergy showed a different contribution of all muscles except for the gluteus maximus. The reorganization of the motor modules provide evidence that the compromised locomotor function is clearly represented by the reassembly of the motor module (Figure 3).

Kargo and colleagues showed *in silico* in 2010 (43) that proprioceptive regulation of muscle synergies can be used to generate meaningful limb trajectories in the frog. Our results provide further *in vivo* evidence that proprioceptive feedback affects both the spatial (motor modules) and temporal (motor primitives) components of muscle synergies. Average differences between the contribution of single muscles within each synergy were bigger in swimming, indicating a higher level of modularity degradation when feedback from both muscle spindles and GTOs is impaired.

### Feedback from muscle spindles regulates adjustments to external perturbations

From the thoughts above, it is clear that the lack of proprioceptive feedback induces alteration in the modular control of locomotion. Yet, the data recorded from the animals of GR1 cannot explain how muscle spindle-deficient mice respond to external perturbations. To investigate how the spinal circuitry adjusts locomotor pattern to accommodate external perturbations, we electrically stimulated the saphenous nerve during walking in a small group (GR2, n = 5) of both WT and *Egr3^−/−^* animals. Stimulation was delivered at regular intervals during locomotion, resulting in a cautious way of walking between two consecutive stimulations. In a previous study on humans, we had healthy young adults walk and run over uneven surfaces (34). What we found was that, despite generally preserving the structure of motor modules (spatial organization), our participants responded to the irregularities of terrain by widening the motor primitives (temporal organization). The widening can be seen as a modification of the state of the system. Therefore, if functionality is maintained (i.e. locomotion can still take place without major interruptions), we can assume that the system flexibly selects slightly different modes of operation (44). A widening in the temporal structure of locomotor patterns increases the fuzziness of the temporal boundaries. This creates a robust “control’s buffer” that might allow an easier shift from one synergy to the next (34, 45). Data recorded from WT mice during cautious walking was interestingly similar. We found a general preservation of the motor modules in both WT and *Egr3^−/−^* mice when comparing normal (no stimulation) and cautious (with stimulation) walking. Only in WT, however, there was a significant widening in the motor primitives in cautious walking, specifically in the first (weight acceptance and propulsion) and third (late swing) synergy. *Egr3^−/−^* mice, on the contrary, did not show any significant widening nor timing shift when we compared normal and cautious walking. We can argue that muscle spindle-deficient animals are not able to adjust their control strategy when a source of external perturbation is introduced. Instead, similarly to what we found in humans walking or running on uneven surfaces (34), WT animals increase the control’s robustness by widening the motor primitives when forced to cope with external perturbations.

### Conclusion

The concept that proprioceptive feedback is an important factor for controlling movement of insects and vertebrates is well established (39, 46–53). However, with this study we provided new proof that the partial or complete absence of proprioceptive signals can affect the accuracy of the modular organization of walking and swimming in mice. We found that, in both walking and swimming, muscle spindle-deficient animals show a temporal rearrangement of the motor primitives’ shape (i.e. loss of accuracy). In addition, we observed a general preservation of the motor modules’ structure in walking and a strong loss in the coordination of the modularity in swimming. Moreover, when electrical stimulation was applied to the saphenous nerve, only WT animals reacted by prolonging the active time of motor primitives, whereas *Egr3^−/−^* mice did not. Given that muscle spindles in *Egr3^−/−^* mice regress immediately after birth (20, 54), we cannot exclude a certain amount of long-term adaptation in the modular control of the locomotor patterns. Nevertheless, we speculate that an acute removal of muscle spindles (8) in the adult mouse could alter the locomotor pattern in an even more amplified fashion.

## Materials and methods

All procedures were performed according to the guidelines of the Canadian Council on Animal Care (CCAC) and approved by the local councils on animal care of Dalhousie University.

### Mouse lines

All recordings were conducted on adult (older than 50 days old) *Egr3* knockout mice (*Egr3^−/−^*) and wild type (WT) mice of either sex. In *Egr3^−/−^* mice, all muscle spindles are ablated with the GTOs left intact (20, 55).

### Surgeries

Each WT (N=21) and *Egr3^−/−^* (N=22) mouse received an electrode implantation surgery as previously described (6). Depending on the animal, a set of three to six bipolar EMG electrodes were implanted (56, 57) to as many muscles of the right hind limb. The muscles that were implanted were chosen among the following: *gluteus maximus* (hip rotator, MA), *iliopsoas* (hip flexor, IP), *vastus lateralis* (knee extensor, VL), *biceps femoris* (anterior, hip extensor and knee flexor, BF), *semitendinosus* (hip extensor and knee flexor, ST), *gastrocnemius lateralis* (ankle extensor, GL) and *tibialis anterior* (ankle flexor, TA). Four WT and five *Egr3^−/−^* mice received an additional nerve stimulation cuff electrode implant (58). Briefly, mice were anesthetized with isoflurane and ophthalmic eye ointment was applied to the eyes. The skin was sterilized by using a three-part skin scrub by means of Hibitane (Chlorhexidine gluconate 4%), alcohol, and povidone-iodine. Small skin incisions were made to the neck region and to the right hind leg to expose the target muscles and the saphenous nerve. The electrodes were then drawn from the neck incision to the leg incisions subcutaneously and the headpiece connector was sutured to the skin around the neck incision. Afterwards, each electrode was implanted to the muscles as described before (56), the incisions were closed, anesthetic was discontinued and buprenorphine (0.03 mg/kg) and ketoprofen (5.00 mg/kg) were injected subcutaneously as analgesics. The cuff electrode was implanted around the saphenous nerve, a nerve carrying cutaneous afferent fibers from the anterior part of the distal hind leg to the spinal cord (59, 60). The mice were then placed in a heated cage for 3 days and finally returned to their regular mouse rack. Food mash and hydrogel was provided for the first 3 days after the surgery. Handling of the mice was avoided until they were fully recovered and at least for one week. Additional injections of buprenorphine were performed in 12-hour intervals for 48 hours.

### EMG and kinematics recordings

After recovery from surgery, the animals were assigned to two experimental setups. One group (henceforth GR1) of 17 WT and 17 mutants completed two different tasks (6). First, they freely walked at a fixed speed of 0.2 m/s on a custom treadmill (workshop of the Zoological Institute, University of Cologne) for a variable amount of time, depending on each animal’s will to walk. After the recordings during walking, 11 of these WT and 11 of these *Egr3^−/−^* mice were placed in a tank filled with water at a temperature of circa 24 °C for around 120 s. A second group (henceforth GR2) of 4 WT and 5 mutants walked on the treadmill at 0.3 m/s while one step was perturbed every second by stimulating the saphenous nerve electrically (five 0.2 ms impulses at 500Hz) that elicited stumbling corrective reaction (SCR). The strength of the stimulation was set to be 1.2 times the current that was necessary to elicit the slightest response in the tibialis anterior muscle during resting, which varied between 96 to 1200 μAmp from animal to animal (8). These frequent SCRs caused the mice to walk cautiously, therefore we called these trials “cautious walking.” Three steps prior to the SCR were taken for data analysis. Only one recording session was performed with each mouse to avoid a learning effect.

The EMG recordings in both groups (GR1 and GR2) were conducted by connecting the electrodes via the headpiece to an amplifier (model MA 102, custom built in the workshop of the Zoological Institute, University of Cologne, DEU) and recorded with a sampling rate of 10 kHz. The animals of GR2 were not implanted with the MA (*gluteus maximus*) electrode, thus only six muscles were recorded. During walking, the kinematics of the hindlimb were acquired through motion capture with a high-speed camera operating at 250 Hz (IL3, Fastec Imaging). Under brief anesthesia with isoflurane, six custom-made cone-shaped reflective markers (1-2 mm in diameter) were placed unilaterally on the limb interested by the EMG recordings. Namely, the markers were attached at the level of the anterior tip of the iliac crest, hip joint, knee joint, ankle joint, the metatarsal-phalangeal joint, and the tip of the fourth digit. Due to the large slippage of the skin above the knee joint, the knee values were calculated by triangulation using the hip and ankle coordinates together with the measured length of femur and tibia (56). Kinematics were post-processed using the motion analysis software Motus (Vicon Motion Systems, GBR) or a custom-made software written by Dr. Nicolas Stifani with ImageJ (KinemaJ) and R (KinemaR) (61, 62). None of the animals were trained for the experiments to avoid the possible complications of different learning capabilities.

### Cycle breakdown

The cycle breakdown was obtained via two different approaches for walking and swimming. In walking, the foot touchdown and lift-off were calculated defining the stance phase as the time interval in which the speed of the toe marker in the anteroposterior direction was matching the speed of the treadmill’s belt (i.e. null derivative, calculated on the raw kinematics). In swimming, the cycles were identified by searching for the onset of the *iliopsoas* muscle activation through visual inspection of the EMG data. Since in 6 out of 17 animals of GR1 the recorded EMG activity of the *iliopsoas* muscle was not clean enough to allow the breakdown of the swimming cycle, these animals were excluded from the analysis. In summary, we analyzed the data coming from 17 WT and 17 *Egr3^−/−^* mice (GR1) during treadmill walking at 0.2 m/s, 11 WT and 11 *Egr3^−/−^* mice (GR1) during free swimming, 4 WT and 5 *Egr3^−/−^* mice (GR2) during treadmill walking at 0.3 m/s with and without electrical stimulation of the saphenous nerve to elicit a stumbling corrective reaction (i.e. what we call “cautious walking”). In the dataset generated from the recordings with electrical stimulation (GR2), the steps in which stimulation occurred were excluded from the analysis. This was done in order to avoid data contamination, particularly concerning EMG.

### Local dynamic stability assessment

In order to assess the local dynamic stability of GR1’s animals during treadmill walking, we used the maximum finite-time Lyapunov exponent (MLE) (24, 63). We calculated the MLEs of three hindlimb joint angles from raw kinematics. The time series describing the evolution of the three angles over time were prepared as follows. For being able to compare data coming from different trials and animals, we subtracted the minimum value from each time series and then normalized the values to the maximum, so that joint angles could only vary between 0 and 1. Then, we linearly interpolated each time series to the same amount of points per gait cycle. To do so, we calculated the average number of recorded kinematic points per gait cycle across all the available trials (23, 64), which was equal to 70. Conventionally, the MLE analysis requires large data sets collected over a big number of gait cycles (23, 24). In our case, however, the minimum number of completed gait cycles across all trials of all animals was only six. Given this limitation, we treated the data in order to increase the number of available cycles using a resampling approach, similar to bootstrapping. The time series for WT and *Egr3^−/−^* treadmill walking (GR1) containing the joint angle values were built as follows. Since the sample size of GR1 was 34 (17 WT and 17 *Egr3^−/−^* animals), for both WT and *Egr3^−/−^* we created 17000 vectors (so 34000 in total) of length 17 containing values from 1 to 17, randomly sampled with replacement (which means that every number could appear more than once). Each number in every vector represented one of the 17 WT or *Egr3^−/−^* animals. Then, to each number we assigned data recorded during 6 consecutive gait cycles, randomly selected from the available cycles recorded for that animal. Then, for the creation of a bootstrapped trial, each 17 sets of 6 cycles were concatenated in a single time series consisting of 17 * 6 = 102 cycles, for a total of *N* = *102* * *70* = *7140* time points. Following this procedure, we then obtained for both WT and *Egr3^−/−^* mice 1000 bootstrapped samples (subsets of the initial 17000) containing 17 bootstrapped trials each for subsequent calculations. By using the minimum number of recorded gait cycles (six), we weighted the contribution of each animal to the data set equally.

Afterwards, we used each new time series to reconstruct the state of the mouse system. The state of a dynamical system is described by a number of variables which is roughly equal to the number of degrees of freedom of that system. The state space is a set of all the possible states of a system at any given time, the variables of which might be position, velocity, temperature, color, species and many others (65, 66). The behavior of a chaotic dynamical system can be reconstructed by a small set of observations on its state (e.g. the hindlimb joint angles in our case) without losing information on its properties. We used the “delay embedding theorem” (67) to reconstruct the state space starting from the three joint angles. The reconstructed time series were obtained as follows:

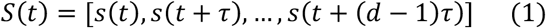

with *S(t)* being a *D* × *d* matrix, where *s(t)* is the input time series (i.e. one of the three joint angles), *d* is the embedding dimension (i.e. the dimension of the state space), *τ* is the time delay and *D* = *N* − (*d − 1*)*τ*. The time delays (*τ*, expressed in number of points in the time series) were calculated for each bootstrapped time series as the first minimum of the Average Mutual Information function (68), implemented using the algorithm of Cellucci (69) with bias correction of Roulston (70). The number of embedding dimensions *d* was extracted through a Global False Nearest Neighbors analysis (71) for each bootstrapped time series, with a threshold of five per cent. In MLE analysis, different values of *τ* and *d* can yield very different state-space reconstructions and therefore lead to different results (72, 73). To ensure the comparability between WT and *Egr3^−/−^*, we took the rounded mean values of *τ* and *d* obtained from all the 17000 state space reconstructions (*τ_hip_* = 20.0 ± 1.1, *τ_knee_* = 15.0 ± 4.2, *τ_ankle_* = 15.0 ± 5.8 data points; *d_hip_* = 5.0 ± 0.4, *d_knee_* = 5.0 ± 0.1, *d_ankle_* = 5.0 ± 0.1). After the reconstruction of the times series, we used the Rosenstein algorithm (74) to calculate each point’s trajectory divergence from the closest (in terms of Euclidean distance) trajectory. The MLE were then calculated as the slope (approximated with a linear fit) of the resulting divergence curve, specifically in the region going from 0 to 70 points (i.e. from 0 to 1 gait cycles (63)). This was done to ensure that the considered part of the curve was not yet saturated, as explained by Rosenstein *et al*. (74). Analysis of the data was performed on MATLAB 2014b (Mathworks Inc., Natick, United States).

### Muscle synergies extraction

Muscle synergies data were extracted through a custom script (R v3.5.1, R Found. for Stat. Comp.) using the classical Gaussian NMF algorithm (19, 21, 22, 34). The code is available in the supplementary dataset S1, together with some example data. The raw EMG signals were band-pass filtered within the acquisition device (cut-off frequencies 10 and 500 Hz). Then the signals were high-pass filtered, full-wave rectified and lastly low-pass filtered using a 4th order IIR Butterworth zero-phase filter with cut-off frequencies 50 Hz (high-pass) and 30 Hz (low-pass for creating the linear envelope of the signal). The amplitude of each EMG recording was normalized to the maximum activation recorded for every individual muscle. Each walking and swimming cycle was time-normalized to 200 points (34), so that all cycles had the same length. In walking, we assigned 100 of the 200 total points to the stance and the remaining 100 points to the swing phase (34). For the analysis we considered the muscles described above in the section “Surgeries” (MA only in GR1, IP, VL, BF, ST, GL, TA). The m = 7 (for animals of GR1, otherwise m = 6 for animals of GR2) time-dependent muscle activity vectors were grouped in a matrix V with dimension m × n (m rows and n columns). The dimension n, which represents the number of normalized time points, was chosen with a bootstrapping strategy as follows. Since not all animals were implanted with the same amount of electrodes, we could not collect the same amount of data for every muscle. Hence, we searched across all animals for all the recordings which were available for each muscle. Finally, we concatenated the available cycles, creating a vector of EMG activations for each muscle containing the data belonging to different animals. For instance, the minimum available number of concatenated cycles recorded from the animals of GR1 was 46 for walking (n = 9200) and 22 for swimming (n = 4400). To create the matrix V containing the recorded EMG activities, we randomly sampled (with replacement) 46 cycles for walking and 22 for swimming for each muscle. The same procedure was adopted with the data recorded from the animals of GR2, where the minimum number of cycles was 43 for normal and 245 for cautious (with electrical stimulation of the saphenous nerve) walking (minimum values across both WT and *Egr3*^−/−^ animals obtained excluding those cycles which were interested by the electrical stimulation, as mentioned above). Then, V was factorized using NMF so that V ≈ V_R_ = WH. The new matrix V_R_, reconstructed multiplying the two matrices W and H, is an approximation of the original matrix V. The motor primitives (14, 19) matrix H contained the time-dependent coefficients of the factorization with dimensions r × n, where the number of rows r represents the minimum number of synergies necessary to satisfactorily reconstruct the original set of signals V. The motor modules (18, 19) matrix W, with dimensions m × r, contained the time-invariant muscle weightings, which describe the relative contribution of single muscles within a specific synergy (a weight was assigned to each muscle for every synergy). H and W described the synergies necessary to accomplish the required task (i.e. walking or swimming). The update rules for W and H are presented in Equation (2) and Equation (3).

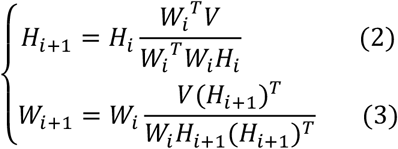

The quality of reconstruction was assessed by measuring the coefficient of determination R^2^ between the original and the reconstructed data (V and V_R_, respectively). The limit of convergence for each synergy was reached when a change in the calculated R^2^ was smaller than the 0.01% in the last 20 iterations (19), meaning that, with that amount of synergies, the signal could not be reconstructed any better. This operation was first completed by setting the number of synergies to 1. Then, it was repeated by increasing the number of synergies each time, until a maximum of five synergies. The number five was chosen to be lower than the number of muscles, since extracting a number of synergies equal to the number of measured EMG activities would not reduce the dimensionality of the data. Specifically, five is the rounded 75% of seven, which is the number of considered muscles. For each synergy, the factorization was repeated 10 times, each time creating new randomized initial matrices W and H, in order to avoid local minima (19). The solution with the highest R^2^ was then selected for each of the five synergies. To choose the minimum number of synergies required to represent the original signals, the curve of R^2^ values versus synergies was fitted using a simple linear regression model, using all five synergies. The mean squared error (19) between the curve and the linear interpolation was then calculated. Afterwards, the first point in the R^2^-vs.-synergies curve was removed and the error between this new curve and its new linear interpolation was calculated. The operation was repeated until only two points were left on the curve or until the mean squared error fell below 10^−4^. This was done in order to search for the most linear part of the R^2^-versus-synergies curve, assuming that in this section the reconstruction quality could not increase considerably when adding more synergies to the model. We extracted muscle synergies in the aforementioned way from 10000 differently re-sampled (with replacement) matrices V, with the idea of building each dataset using data coming from different combinations of animals. This allowed us to obtain 10000 sets of motor modules (W) and motor primitives (H) for subsequent analysis.

By means of the above procedure, we extracted fundamental and combined synergies from the raw EMG data (19). A fundamental synergy can be defined as an activation pattern whose motor primitive shows a single peak of activation (34). When two or more fundamental synergies are blended into one, a combined synergy appears. Due to the lack of consent in the literature on how to interpret them, combined synergies were excluded from the analysis. We compared motor primitives by evaluating the coefficient of determination (R^2^, a metric we also employed, as described above, to estimate the similarity between the original and the reconstructed EMG data after factorization (34)), the center of activity (CoA) and the full width at half maximum (FWHM). We previously found in humans that typical R^2^ values between intraday measurements are, on average, bigger than 0.74 for motor primitives (34). This means that, when measuring the same participant in the same condition twice without replacing the electrodes, the R^2^ similarity between the primitives extracted from the two trials are not likely to give values lower than 0.74. The CoA was defined as the angle of the vector (in polar coordinates) that points to the center of mass of that circular distribution (34, 41, 42). The polar direction represented the cycle’s phase, with angle 0 ≤ θ_t_ ≤ 2π. The following equations define the CoA:

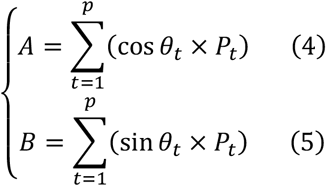

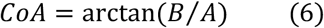

where p is the number of points of each cycle (p = 200) and P is the activation vector. The FWHM was calculated as the number of points exceeding each cycle’s half maximum, after subtracting the cycle’s minimum (34). The CoA and FWHM were analyzed cycle-by-cycle and then averaged.

### Statistics

To evaluate the differences in locomotion parameters between WT and *Egr3^−/−^* animals of GR1, we used a t-test for independent samples or a Wilcoxon signed-rank test for independent samples in case the normality assumptions were not satisfied. The homogeneity of variances was tested using the Levene’s test. Significance Level was set to α = 0.05. The same approach was applied for evaluating the differences between walking with and without electrical stimulation in GR2.

For both GR1 and GR2, we estimated the 95% confidence interval of the bootstrapped FWHM and CoA values for motor primitives using the 2.5% sample quantile as the lower bound and the 97.5% sample quantile as the upper bound (R v3.5.1, R Found. for Stat. Comp.). The same procedure was employed to compare motor modules, where confidence intervals were calculated based on the muscle-by-muscle differences between conditions.

For the MLE data, which referred solely to the data acquired from the animals of GR1 during walking (17 WT and 17 *Egr3^−/−^* animals), we calculated the difference of the means between *Egr3^−/−^* and WT 1000 times (bootstrap, resampling with replacement) for an empirical distribution. The 95% confidence interval for the expectation value of difference was estimated by using the 2.5% sample quantile as the lower bound and the 97.5% sample quantile as the upper bound of this sample of differences (n = 1000). Calculations were done with MATLAB 2014b (Mathworks Inc., Natick, United States). Moreover, we calculated the effect size Hedges’ g as an unbiased estimator for the standardized mean differences between independent groups. The approximate distribution of the effect size g was calculated from the bootstrapped MLE sample pairs (WT and *Egr3^−/−^* mice, 1000 bootstrapped samples of 17 elements each). Confidence intervals were taken from this distribution as described above.

## Acknowledgments

We thank W. Tourtellotte for providing the breeders, from which our colony was established and Brenda Ross for the technical assistance with, surgeries and experiments and maintaining the mouse colony. This work was supported by the Dalhousie Medical Research Foundation and Natural Sciences and Engineering Research Council (NSERC).

## Supplementary Information

In this study, we made use of non-negative matrix factorization (NMF) to extract muscle synergies from electromyographic (EMG) data. We implemented the NMF algorithm in R version 3.5.1 (R Foundation for Statistical Computing, R Core Team, Vienna, Austria), a programming language available in a free software environment. However, even if the software does not require a paid license, often researchers are either not confident with or prefer not to spend time writing the code required to perform NMF. We make available, as we recently did with human data (1), an example open access data set of EMG and muscle synergy data for murine walking and swimming. The data presented in this supplementary information part is available in three formats: 1) the raw EMG of two example trials (one recorded during walking and the other during swimming in a wild type animal, six muscles), unprocessed together with the touchdown and lift-off timings of the recorded limb for walking and the cycle timings for swimming; 2) the filtered and time-normalized EMG and 3) the muscle synergies extracted via NMF. Moreover, we provide the R code for obtaining the results described in the previous three points.

### Description of the data

We do not report any metadata, since trials are relative to a single representative animal. The R code is profusely commented.

#### Gait cycle parameters

The files containing the walking and swimming cycle breakdown are available in ASCII and RData (R Found. for Stat. Comp.) format. The files are structured as data frames with as many rows as the number of available cycles and two columns. The first column of the walking trial contains the touchdown incremental times in seconds. The second column contains the duration of each stance phase in seconds. The only column of the swimming trial contains the starting times of each cycle. Each trial is saved both as a single ASCII file and as an element of a single R list. Trials are named like “CYCLE_TIMES_A0001_W”, where the characters “CYCLE_TIMES” indicate that the trial contains the walking or swimming cycle breakdown times, the characters “A0001” indicate the animal number (in this example the first) and the last character indicates the type of locomotion (“W” for walking and “S” for swimming).

#### EMG data

The files containing the raw, filtered and the normalized EMG data are available in ASCII and RData (R Found. for Stat. Comp.) format. The raw EMG files are structured as data frames with as many rows as the recorded data points and seven columns. The first column contains the incremental time in seconds. The remaining six columns contain the raw EMG data, named with muscle abbreviations that follow those reported in the methods section of this paper (note that in these examples, only six muscles are presented, see methods). Each trial is saved both as a single ASCII file and as an element of a single R list. Trials are named like “RAW_EMG_A0001_W”, where the characters “RAW_EMG” indicate that the trial contains raw emg data, the characters “A0001” indicate the animal number (in this example the first) and the last character indicates the type of locomotion (“W” for walking and “S” for swimming). The filtered and time-normalized emg data is named, following the same rules, like “FILT_EMG_A0001_W”.

#### Muscle synergies

The files containing the muscle synergies extracted from the filtered and normalized EMG data are available in ASCII and RData (R Found. for Stat. Comp.) format. The muscle synergies files are divided in motor primitives and motor modules and are presented as direct output of the factorization and not in any functional order.

Motor primitives are data frames with a number of rows equal to the number of synergies (which might differ from trial to trial) and a number of columns equal to the number of cycles times 200 points (see methods). The rows contain the time-dependent coefficients (motor primitives), one row for each synergy (named e.g. “Syn1, Syn2, Syn3”, where “Syn” is the abbreviation for “synergy”). Each gait cycle contains 200 data points, which in walking are divided in 100 for the stance and 100 for the swing phase. Each set of motor primitives relative to one synergy is saved both as a single ASCII file and as an element of a single R list. Trials are named like “SYNS_H_P0026_02”, where the characters “SYNS_H” indicate that the trial contains motor primitive data, the characters “A0001” indicate the animal number (in this example the first) and the last character indicates the type of locomotion (“W” for walking and “S” for swimming). Motor modules are data frames with as many rows as the number of considered muscles (seven in this example) and a number of columns equal to the number of synergies (which might differ from trial to trial). The rows, named with muscle abbreviations that follow those reported in the methods section of this paper, contain the time-independent coefficients (motor modules), one for each synergy and for each muscle. Each set of motor modules relative to one synergy is saved both as a single ASCII file and as an element of a single R list. Trials are named like “SYNS_W_A0001_W”, where the characters “SYNS_W” indicate that the trial contains motor module data, the characters “A0001” indicate the animal number (in this example the first) and the last character indicates the type of locomotion (“W” for walking and “S” for swimming).

Since they are time-invariant coefficients, motor modules are usually represented with bar graphs. On the contrary, motor primitives describe the evolution over time of the basic activation patterns and are therefore better represented with time-dependent curves. When multiplying and summing synergy-by-synergy the elements of the two matrices W (motor modules) and H (motor primitives), it is possible to reconstruct the original set of EMG data.

#### Code

All the code used for the preprocessing of EMG data and the extraction of muscle synergies is available in R (R Found. for Stat. Comp.) format. Explanatory comments are profusely present throughout the scripts (“SYNS.R”, which is the main script and “fun_synsNMFn.R”, which contains the NMF function).

### Dataset S1 (separate file)

Dataset S1 contains data from two example trials, one recorded during walking and the second during swimming in a wild type animal. The raw and the filtered and time-normalized EMG activities are available in both ASCII and RData (R Found. for Stat. Comp.) format. The same goes for the synergy data and the cycle times (useful to normalize the EMG data). All the R code that can be used to extract synergies from this kind of EMG data is included as well.

